# Comparisons of the antibody repertoires of a humanized rodent and humans by high throughput sequencing

**DOI:** 10.1101/776385

**Authors:** Collin Joyce, Dennis R. Burton, Bryan Briney

## Abstract

The humanization of animal model immune systems by genetic engineering has shown great promise for antibody discovery, tolerance studies and for the evaluation of vaccines. Assessment of the baseline antibody repertoire of unimmunized model animals will be useful as a benchmark for future immunization experiments. We characterized the heavy chain and kappa chain antibody repertoires of a model animal, the OmniRat, by high throughput antibody sequencing and made use of two novel datasets for comparison to human repertoires. Intra-animal and inter-animal repertoire comparisons reveal a high level of conservation in antibody diversity between the lymph node and spleen and between members of the species. Multiple differences were found in both the heavy and kappa chain repertoires between OmniRats and humans including gene segment usage, CDR3 length distributions, class switch recombination, somatic hypermutation levels and in features of V(D)J recombination. The Inference and Generation of Repertoires (IGoR) software tool was used to model recombination in VH regions which allowed for the quantification of some of these differences. Diversity estimates of the OmniRat heavy chain repertoires almost reached that of humans, around two orders of magnitude less. Despite variation between the species repertoires, a high frequency of OmniRat clonotypes were also found in the human repertoire. These data give insights into the development and selection of humanized animal antibodies and provide actionable information for use in vaccine studies.

## Introduction

A major challenge in human vaccine science is finding appropriate models for studying antibody responses. Animals such as mice, rabbits and monkeys have typically been used in the past and the small animals, in particular, have been favored for ease of immunization, cost reasons and the ability to extensively biopsy post-immunization. One limitation is their use of non-human immunoglobulin (V, D, J) genes in antibodies which can be restricted in their specificity^1^, and/or lack residues needed for priming by a germline targeting immunogen^2^. One approach to solving this problem of wild-type animal models is to use humanized immunoglobulin loci-transgenic rodents^3, 4^. The first demonstration of a transgenic rodent with the ability to express human IgM was 30 years ago^5^. Since then, advances in genetic engineering technologies allowed for the first transgenic mice strains that express fully human antibodies^6, 7^. Today, many new transgenic animal models have been developed including rodents, chickens, rabbits and cows^8^. These animal models have been used extensively for the discovery of monoclonal antibodies (mAbs)^9^, tolerance studies^10^ and more recently for modelling human antibody responses to vaccine candidates^3, 4^. Here we focus on one such animal model: a rat with expression of humanized chimeric antibodies.

The generation of antibody diversity begins with the development of B cells in the bone marrow. Three unlinked loci contain the immunoglobulin gene segments necessary for the assembly of an antibody: one heavy chain locus on chromosome 14 and two light chain loci (lambda and kappa found on chromosomes 2 and 22 respectively). Large pre-B cells derived from common lymphoid progenitors randomly join VH, DH, and JH gene segments to produce a heavy chain. This process requires V(D)J recombinase: a protein complex that contains RAG1, RAG2 and Artemis (among others). P and N nucleotides are added in the VH-DH and DH-JH junctions by Artemis and Tdt, dramatically increasing sequence diversity. After successful pairing of this newly formed heavy chain with surrogate light chain (SLC), recombination of a light chain from V and J gene segments of the kappa or lambda loci occurs and the B cell swaps the SLC for this new light chain. Unless the immature B cell is autoreactive or anergic and undergoes receptor editing or clonal deletion, it matures into a naïve B cell and migrates to the periphery whereupon it can become activated by encountering antigen and form germinal centers with help from T-cells. Sequence diversity is again enhanced in the germinal center by somatic hypermutation (SHM) and/or class switch recombination (CSR), two processes that depend on activation induced cytidine deaminase (AID). The OmniRat was created by genomic integration of human immunoglobulin (Ig) loci on a background of inactivated endogenous rat Ig loci. It expresses chimeric heavy chains and fully human light chains^11, 12^. We sought to characterize the circulating antibody repertoire diversity in this animal and make comparisons to humans.

High throughput antibody sequencing has been used to describe the circulating antibody repertoire of organisms, including more recently at unprecedented depth in humans^13^. Reverse transcription of antibody RNA and combined tagging with unique molecular identifiers (UMIs) have allowed us^14^ and others^15, 16^ to correct for error and bias in antibody sequencing. Using these methods to gain insight into the antibody repertoire of OmniRats, we ask whether or not it accurately represents that of humans, and by extension allows for usefulness in the approximation of the human antibody response. We postulate that there are major differences in the repertoires due to distinctness in the Ig loci genomic structure and genes that shape antibody diversity between species. Here, we provide the most thorough description of humanized transgenic rodent antibody repertoires to date and leverage a novel extremely deep human dataset to make comparisons with implications of immediate use as a reference for OmniRat immunization studies.

## Results and Discussion

We individually separated total RNA from spleens and lymph nodes of three unimmunized OmniRats and PCR amplified the heavy and kappa chain antibody V gene segments. The resulting amplicons were subjected to high throughput sequencing in conjunction with preprocessing and annotation by the AbStar analysis pipeline^17^ (Methods) which resulted in a mean of ∼3 × 10^6^ processed heavy chain sequences and ∼1.5 × 10^6^ processed kappa chain sequences per transgenic animal (Table S1). Two previously published datasets^13, 18^ of the same 10 humans which together contain a mean of ∼3.6 × 10^7^ processed heavy chain sequences and ∼1.5 × 10^6^ processed kappa chain sequences per individual were used for comparison.

### Gene usage comparisons between different tissue sources, between individual OmniRats and between OmniRats and humans

We started by making intra-animal comparisons, intra-species comparisons and inter-species comparisons of the immunoglobulin gene segment usage frequencies for each antibody repertoire by performing hierarchical clustering (Figure 1) and linear regression analysis (Figures S1 and S2). Repertoires were found to cluster by species and tissue when variable heavy (VH) (Figure 1a), diversity heavy (DH) (Figure 1b), joining heavy (JH) (Figure 1c) and variable kappa (VK) (Figure S5a), but not joining kappa (JK) (Figure S5b) gene usage was examined. Differences between the lymph node and spleens of individual OmniRats were next investigated. VH gene, DH gene and JH gene usage frequencies between these tissues were highly correlated (Figure S1), although a few VH gene segments were overrepresented in spleen as compared to lymph nodes including VH4-34, VH4-59, VH5-1 among others (Figure 1a, Figure S3a). DH gene and JH gene usage remained highly correlated with minor differences in specific genes (Figures 1b-c, Figure S1, Figures S3b-c). Inter-animal spleen gene usage was highly correlated for all three heavy chain gene segments (Figure S2). Inter-human comparisons yielded similar, albeit slightly less correlated results (Figure S2). Intra-species VH and DH usage comparisons made show no correlation with lower R-squared values than any other previous comparison, while surprisingly JH gene usage was highly correlated (Figure S2). OmniRats show a preference for DH gene families of shorter average length such as DH1, DH5, DH6 and DH7 as compared to humans which show a higher representation of longer DH genes from DH2, DH3, and DH4 families with the exception of DH3-9 which appears at similar frequencies between each species (Figure 1b, Figure S3b). VK and JK gene usage frequencies were very similar for all comparisons made (Figures S1-3, Figure S6).

**Figure 1.**
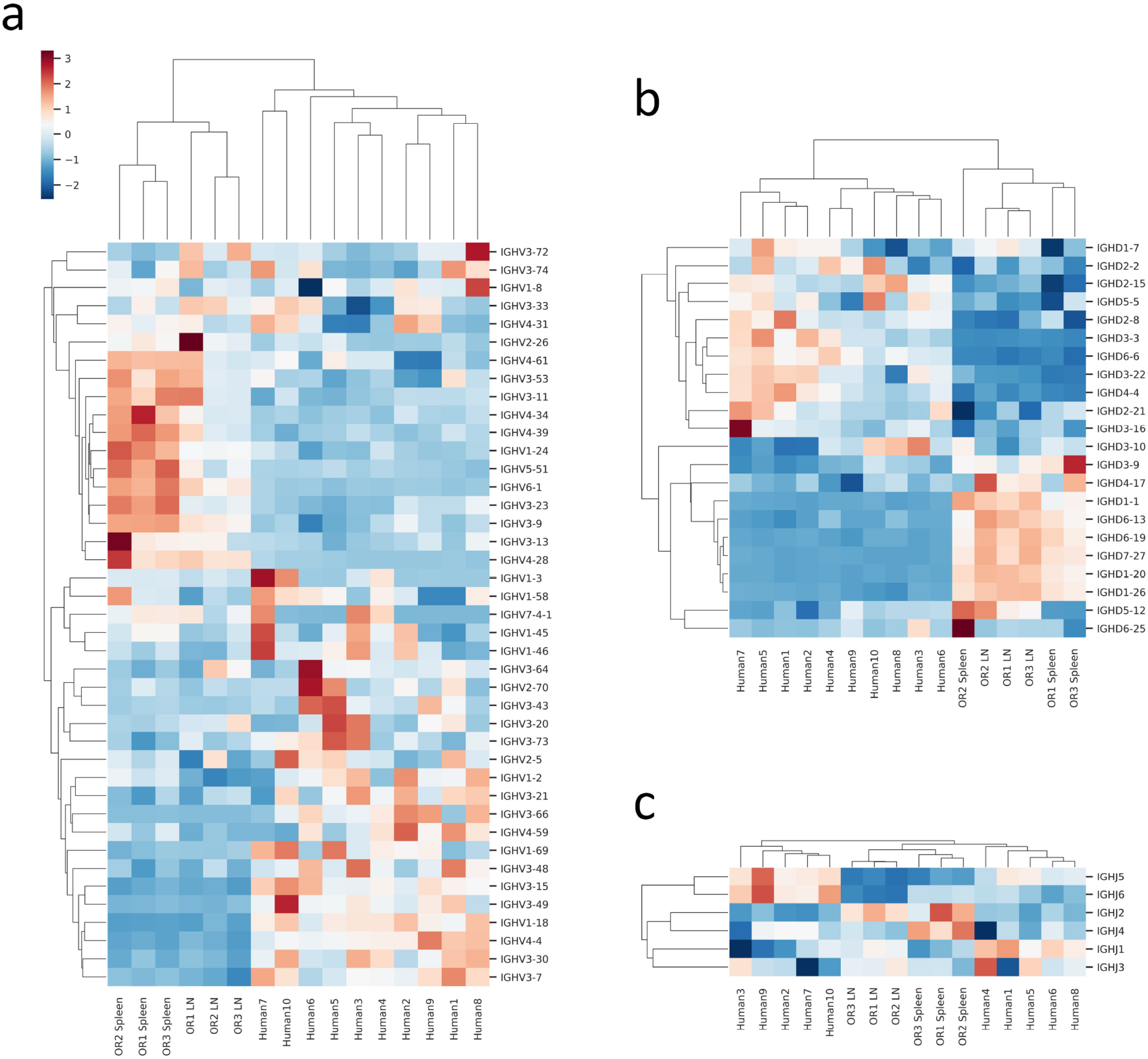
Antibody repertoires cluster by heavy chain gene segment usage. Heatmaps of variable heavy (VH) gene usage in (a), diversity heavy (DH) gene usage in (b) and joining heavy (JH) gene usage in (c). Columns are antibody repertoires and rows are gene segments. Data was scaled by calculating the Z-score for each gene (row) and hierarchical clustering (Euclidean distance metric) was done. A dendrogram representation of clustering is shown and indicates uniqueness in gene segment usage between the lymph node and spleen repertoires of the OmniRat and between the OmniRat and human repertoires. Red and blue indicate high and low Z-scores respectively (legend shown in a), and since it is calculated per gene it represents differences between repertoires and not the relative frequencies of gene segment usage in each repertoire.

### CDR3 comparisons between OmniRats and humans

Differences in CDR3 length distributions of each repertoire were next determined. The mean heavy chain CDR3 (CDRH3) length in humans is 14.8 amino acids, while in the OmniRat we observed a mean CDRH3 length that is shorter with a mean length of 12.1 amino acids (Figure 2a). There are minor differences in the kappa light chain (CDRL3) lengths between species with near identical average lengths of 9.0 and 9.1 for OmniRats and humans respectively (Figure S6c). The frequency of light chains with a CDR3 of 5 amino acids in length is an important consideration when choosing a model animal for vaccination experiments involving the germline targeting immunogen eOD-GT8 which is in human clinical trials^3, 14, 19^. This frequency of 5-amino acid CDRL3s was lower in OmniRats (0.02%) than in humans (0.56%) i.e. a factor of 28 (Figure S6c).

**Figure 2.**
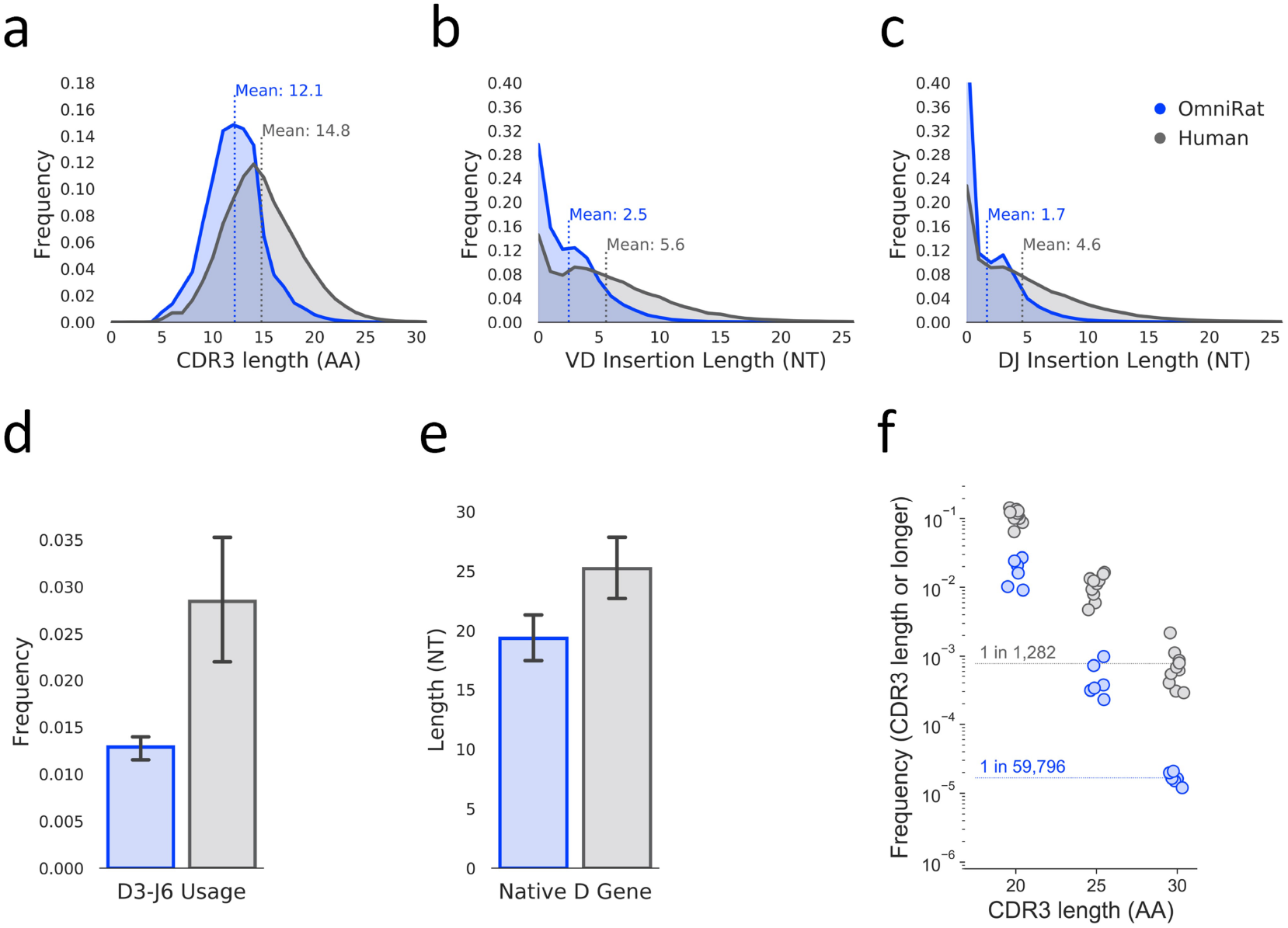
CDRH3 length differences between OmniRat and human repertoires. a) CDRH3 length distribution for each species. Species are colored as in (c). CDRH3 lengths were determined using the ImMunoGeneTics (IMGT) numbering scheme. b) VD insertion length distributions for each species. Species are colored as in (c). c) DJ insertion length distributions for each species. Species are colored as in (c). d) Average frequency of antibody sequences that use DH family 3 and JH family 6 for each species. Species are colored as in (c). Error bars indicate standard error of the mean. d) Average DH gene segment length (NT) for wild-type (WT) rat (*Rattus norvegicus*) in blue and Homo sapiens in gray. Lengths are calculated by counting the number of nucleotides from DH gene segments from the IMGT gene database for each species. Error bars indicate standard error of the mean. Species are colored as in (c). f) Usage frequency of antibody sequences with the given CDRH3 length or longer for each species. Species are colored as in (c).

After observing a tendency for shorter CDRH3 lengths in the OmniRat as compared to humans, we wanted to know if the number of N and P nucleotide additions in the heavy chain V-D and D-J junction sites were different. Figures 2b and 2c shows average V-D and D-J junction nucleotide addition lengths in the OmniRat are indeed shorter as compared to humans. Nucleotide additions in the V-J junctions of kappa chains are also shorter on average as compared to humans (Figure S6e).

The longest DH gene segments are found in the DH3 family and the longest JH gene segments come from the JH6 gene family. Gene segments from these families are important contributors to the generation of unusually long CDRH3s in humans and are consistently found in certain broadly neutralizing antibodies (bnAbs) that bind to the human immunodeficiency virus (HIV) envelope glycoprotein (Env) protein, indicating the importance of these rearrangements in HIV vaccine studies^20^. On average, the frequency of antibodies with D3-J6 rearrangements in OmniRats is 0.012 with little variation, while in humans the frequency of these antibody species is more variable between subjects with a higher mean of 0.028 (Figure 2d). The preference of OmniRats for shorter CDRH3 lengths and DH gene segments can be placed in the context of shorter DH gene lengths in the wild-type rat (*Rattus norvegicus*) as compared to human DH genes (Figure 2e), indicating a possible biologically intrinsic bias.

We used IGoR^21^ to infer recombination models for each individual repertoire from 100,000 unmutated sequences allowing for the quantification of differences in features of heavy chain VDJ recombination and generated 1,000,000 synthetic sequences per model. CDRH3 length, VD insertion length and DJ insertion length distributions from the synthetic sequence data (Figure S4a-c) were found to be very similar to the observed data (Figure 2a-c). Kullback–Leibler (KL) divergence is a measure of how different two probability distributions are. KL divergence between models (Figure S4d) and model ‘events’ (Figure S4e) were computed as previously described^13^. KL divergence between pairs of OmniRat models was found to be lower than KL divergence between pairs of human models for both complete and all event level calculations. The average pairwise OmniRat model versus human model complete KL divergence calculation was found to be much greater than that of pairwise inter-animal calculations and more than twice that of pairwise inter-human calculations. “D-Gene”, “V-gene trim (3’)”, and “D-gene trim (3’)” were among the events computed to have the mean highest KL divergence from pairwise inter-species event level model comparisons.

### Class switch recombination and somatic hypermutation in OmniRats

In supplementary figure S5a, the frequency of antibody isotypes is shown. The human repertoire contains average frequencies of 0.84 and 0.16 for IgM and IgG respectively as previously published^13^, while in the OmniRat antibody repertoire we observe mean frequencies of 0.15 and 0.003 for lymph node and spleen IgG respectively and means of 0.85 and 0.997 for lymph node and spleen IgM respectively. Mean numbers of variable gene mutations in IgM (Figure S5b), IgG (Figure S5c) and kappa (Figure S6d) sequences of the OmniRat were about half of those found in the human repertoire. The observed increase in SHM of class-switched IgG sequences as compared to IgM sequences in the OmniRat demonstrates the ability of the animal to generate memory B cells.

### Heavy chain diversity estimates in OmniRats

We first examined clonotype (defined as identical VH gene, JH gene and CDRH3 amino acid sequence) diversity of the heavy chain repertoire for each individual animal. All sequences from lymph nodes and spleens were collapsed into unique clonotypes, separately for each tissue and animal. Any clonotype found in both tissue compartments must have originated from different B cells, allowing for the measurement of repeatedly observed clonotypes. Rarefaction curves for each animal were generated (Figure 3a) and indicate a low frequency of repeatedly observed clonotypes. We estimated diversity using Chao 2 (Chao, 1987) and Recon (Kaplinksy, 2016) as previously described^13^. Diversity estimates were similar between the two estimators, (7.6 × 10^6^ - 1.2 × 10^7^) for Chao and (9.4 × 10^6^ - 1.9 × 10^7^) for Recon (Figure 3b). We next estimated heavy chain sequence diversity for each animal (Figure 3c) and again found that both estimators broadly agreed, giving similar values of (5.4 × 10^7^ - 1.0 × 10^8^) for Chao and (8.1 × 10^7^ - 1.3 × 10^8^) for Recon. Previously published estimates of both clonotype and sequence diversity in individual humans^17^ only exceed that in the OmniRats by a maximum two orders of magnitude. This is surprising given that the OmniRat is more restricted in CDRH3 length.

**Figure 3.**
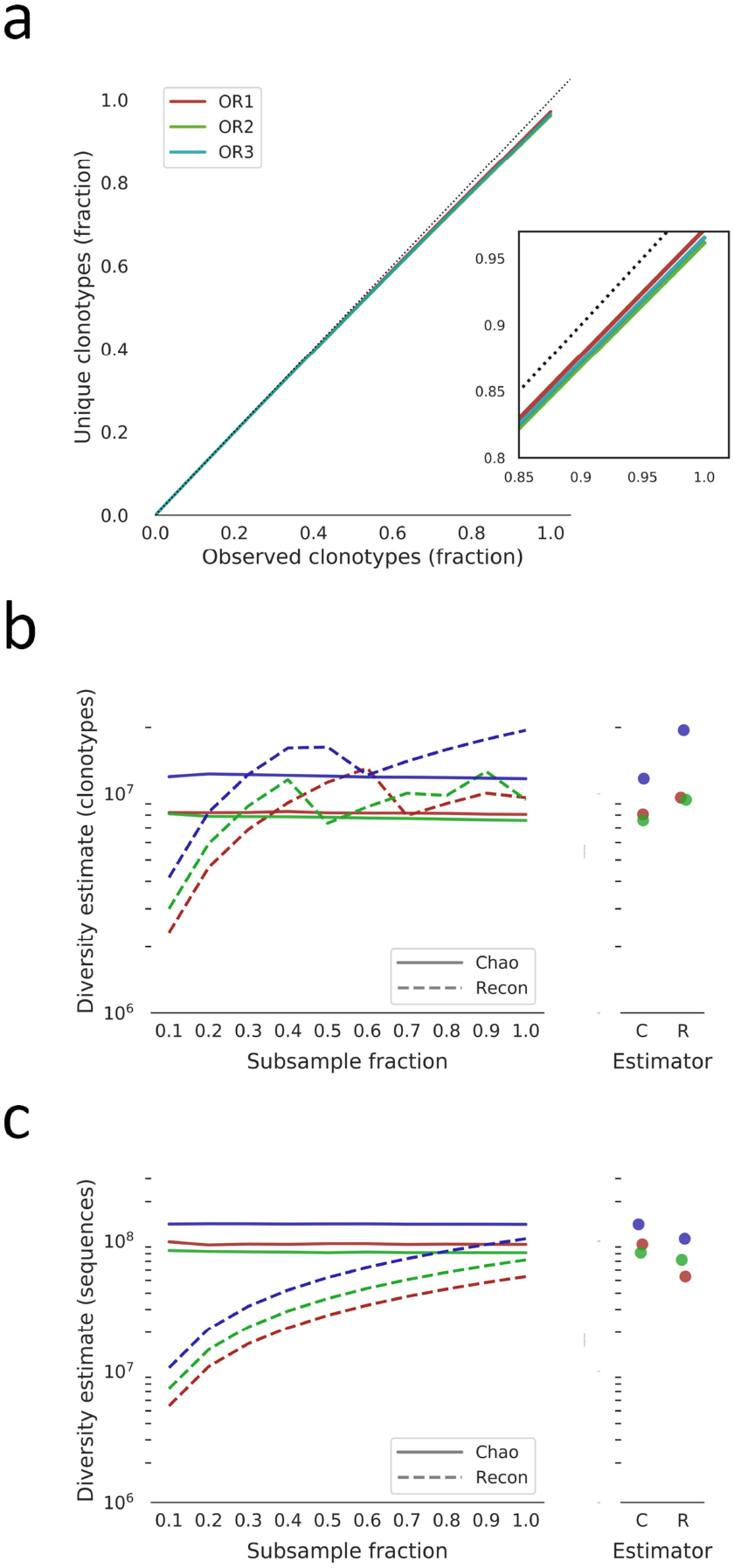
Clonotype and sequence diversity of the OmniRat antibody heavy chain repertoire. a) Clonotype rarefaction curves for each animal. Lines represent the mean of 10 independent samplings, except for the 1.0 fraction which was sampled once. The dashed line indicated a perfectly diverse sample. Inset is a close-up of the rarefaction curve ends. b) Total clonotype repertoire diversity estimates were computed for increasingly large fractions of each animal’s clonotype repertoire. Lines represent the mean of 10 independent samplings, except for the 1.0 fraction which was sampled once. Chao estimates are shown in solid lines and Recon estimates are shown in dashed lines. Lines are colored as in (a). Maximum diversity estimates (from the 1.0 fraction) for each animal is shown in the panel on the right. b) Total sequence repertoire diversity estimates were computed for increasingly large fractions of each animal’s sequence repertoire. Lines represent the mean of 10 independent samplings, except for the 1.0 fraction which was sampled once. Chao estimates are shown in solid lines and Recon estimates are shown in dashed lines. Lines are colored as in (a). Maximum diversity estimates (from the 1.0 fraction) for each animal is shown in the panel on the right.

### Sharing of repertoires between individual OmniRats and between OmniRats and humans

For each combination of two or more animals, we computed the frequency of shared unique heavy chain clonotypes (Fig. 4a). There was on average 9.32% of clonotypes shared between each combination of two OmniRats. Surprisingly, we found that 4.90% of clonotypes were shared between all three of the animals.

**Figure 4.**
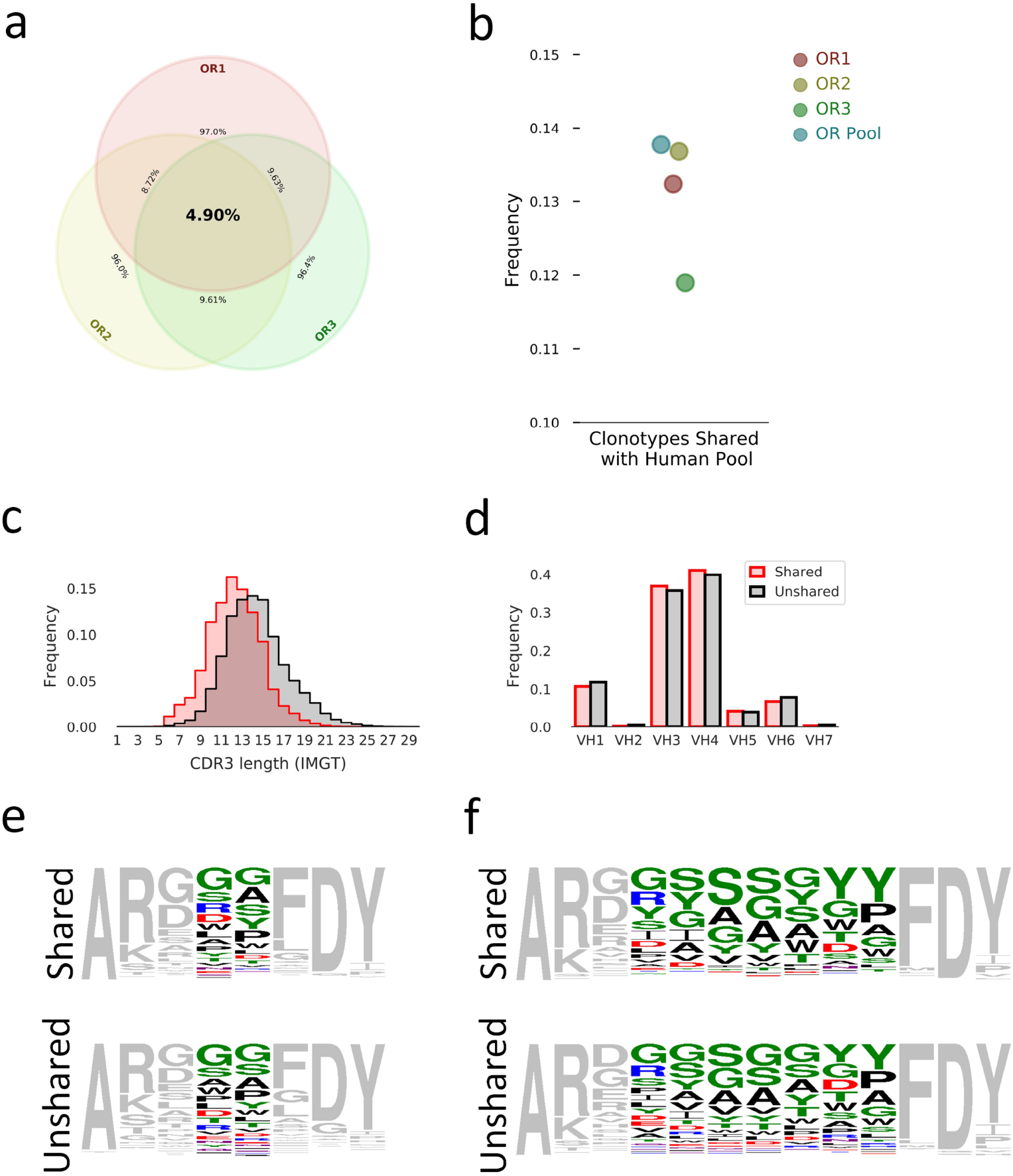
Sharing of clonotypes between repertoires. a) Venn diagram of shared clonotype frequencies between animals. b) Shared clonotype frequencies from individual animals and total animal pool (OR Pool) with total human unique clonotype pool. c) Distribution of CDRH3 length for unshared clonotypes from the animal pool (black) or clonotypes shared by both species pool (red). Distributions are colored as in (d). d) Distribution of VH gene family usage for unshared clonotypes from the animal pool (black) or clonotypes shared by both species pool (red). e), f), Sequence logos of the CDRH3s encoded by shared and unshared clonotypes of length 8 (e) or 13 (f). Head-region amino acid colouring: polar amino acids (GSTYCQN) are green; basic amino acids (KRH) are blue; acidic amino acids (DE) are red; and hydrophobic amino acids (AVLIPWFM) are black. All torso residues are grey

Next, we pooled unique heavy chain clonotypes from all ten human subjects and measured the percentage of clonotypes in each animal that could be found in the total human pool (Figure 4b). We found that (11.9% - 13.7%) of each OmniRat clonotype repertoire and 13.8% of clonotypes combined from all animals could be found in the total human clonotype pool. Shared clonotypes have shorter CDRH3 lengths than unshared clonotypes on average (Figure 4c) which is expected given that sequence diversity is expected to increase as the number of amino acids increases giving less of a chance for sharing. VH gene family usage between shared and unshared clonotypes indicates no major differences (Figure 4d). Sequence logos for 8 amino acid long (Figure 4e) and 13 amino acid long (Figure 4e) CDHR3s from both shared and unshared fractions were made and indicate broad similarity between the two fractions.

## Conclusion

We set out to determine commonalities and differences between OmniRat and human antibody repertoires to be used as a reference for vaccine studies. Our results show that there exists substantial variation in gene usage frequencies and elements of recombination, indicating specific limitations of this animal model for predicting the human immune response. We found that by performing hierarchical clustering on gene segment usage, repertoires clustered together by both species and tissue. Differences in gene segment usage between transgenic animal models and humans, as well as between tissues are expected. For example, multiple human Ig loci transgenic rodents are reported to have gene usage profiles that slightly vary from that of humans^22, 23^. Furthermore, antibody repertoires from separate human tissues are known to deviate strongly enough to be clustered by hierarchical clustering^24^. In our case, the lymph node repertoire from the OmniRat was most likely able to be distinguished from that of the spleen due to the increased presence of antigen-experienced B cells in the latter as shown by somatic hypermutation and class-switched transcript frequencies. This indicates that tissue selection will affect the outcome of an antibody discovery campaign and reinforces evidence for normal B cell development by suggesting the existence of affinity maturation.

Investigation into CDR3 length distributions revealed that the OmniRat prefers shorter CDRH3s as compared to humans. Interestingly, the mechanisms of this preference are due to decreased N additions, and a tendency to incorporate shorter DH gene segments. This result has also been seen in multiple other transgenic and wild-type rodents^22, 23, 25, 26^. The specific reasons remain unclear, although the observation that wild-type rat germline D gene segment lengths are shorter suggests intrinsic species-specific mechanisms of selection as well as differences in Tdt expression during bone marrow B cell development.

The diversity of the OmniRat heavy chain repertoire was shown to be slightly lower than that in humans. Our results indicate biased gene usage and decreased junctional diversity are the primary reasons for the resulting repertoire diversity estimate comparisons. We also showed that there is a much higher frequency of ‘public clonotypes’ or clonotypes shared between members of this species than previously reported in humans^13, 27^. Lower sequence diversity combined with identical genetic background and highly similar gene usage are possible reasons for this result.

In summary, we have determined specific differences between the OmniRat and human antibody repertoires which must be taken into careful consideration when evaluating an antibody response in order to make predictions for human subjects. We have also shown that this animal’s antibodies show signs of class switch recombination, somatic hypermutation and large diversity supporting its value for the discovery of monoclonal antibodies to targets that may not be immunogenic in other models. Even though a high degree of variation exists, we still found many clonotypes to be shared between the species pools. Finally, more studies will need to be done in order to characterize OmniRat serum and memory B cell responses to immunogens.

## Methods

### Next-generation sequencing of OmniRat antibody repertoires

Total RNA from spleens and lymph nodes was extracted (RNeasy Maxi Kit, Qiagen) from each unimmunized transgenic rat (OmniRat, Open Monoclonal Technology Inc., Palo Alto, CA, USA) and antibody sequences were amplified as previously described^13^ except for different primers used during reverse transcription (Table S2). Correct PCR product sizes were verified on an agarose gel (E-Gel EX; Invitrogen) and quantified with fluorometry (Qubit; Life Technologies), pooled at approximately equimolar concentrations and each sample pool was re-quantified before sequencing on an Illumina MiSeq (MiSeq Reagent Kit v3, 600-cycle). All animal experiments were conducted in accordance with the Institutional Animal Care and Use Committee of Scripps Research and approved by the Institutional Research Boards of Scripps Research.

### Processing of next-generation sequencing data

The AbStar analysis pipeline^17^ was used as previously described to quality trim, remove adapters and merge paired sequences. Sequences were then annotated with AbStar in combination with UMI based error correction by AbCorrect (https://github.com/briney/abtools/blob/master/docs/source/abcorrect.rst). Resulting annotated consensus sequences were deposited to MongoDB and Spark database for querying and data analysis in python on Jupyter and Zeppelin Notebooks. For comparisons of frequencies, read counts were scaled for each repertoire as previously described^28^ due to the large differences in the number of reads between species and number of genes that each species expresses.

## Data and Code availability

Sequencing data and code for figures and analysis will be available at www.github.com/CollinJ0/omnirat_paper upon publication of the manuscript.

## Acknowledgements

The authors thank all the study subjects for their participation. This work was supported by the National Institute of Allergy and Infectious Diseases (Consortium for HIV/AIDS Vaccine Development, UM1AI144462 (D.R.B.); Center for Viral Systems Biology, U19AI135995 (B.B.)), the International AIDS Vaccine Initiative (IAVI) through the Neutralizing Antibody Consortium, SFP1849 (D.R.B.), and the Ragon Institute of MGH, MIT and Harvard (D.R.B.).

## Author contributions statement

C.J., D.R.B., and B.B. planned and designed the experiments. C.J. performed experiments. C.J. analysed data. C.J. wrote the manuscript. All authors contributed to manuscript revisions.

## Additional information

### Corresponding authors

Correspondence to Dennis R. Burton or Bryan Briney

---

**Supplementary Figure 1.**
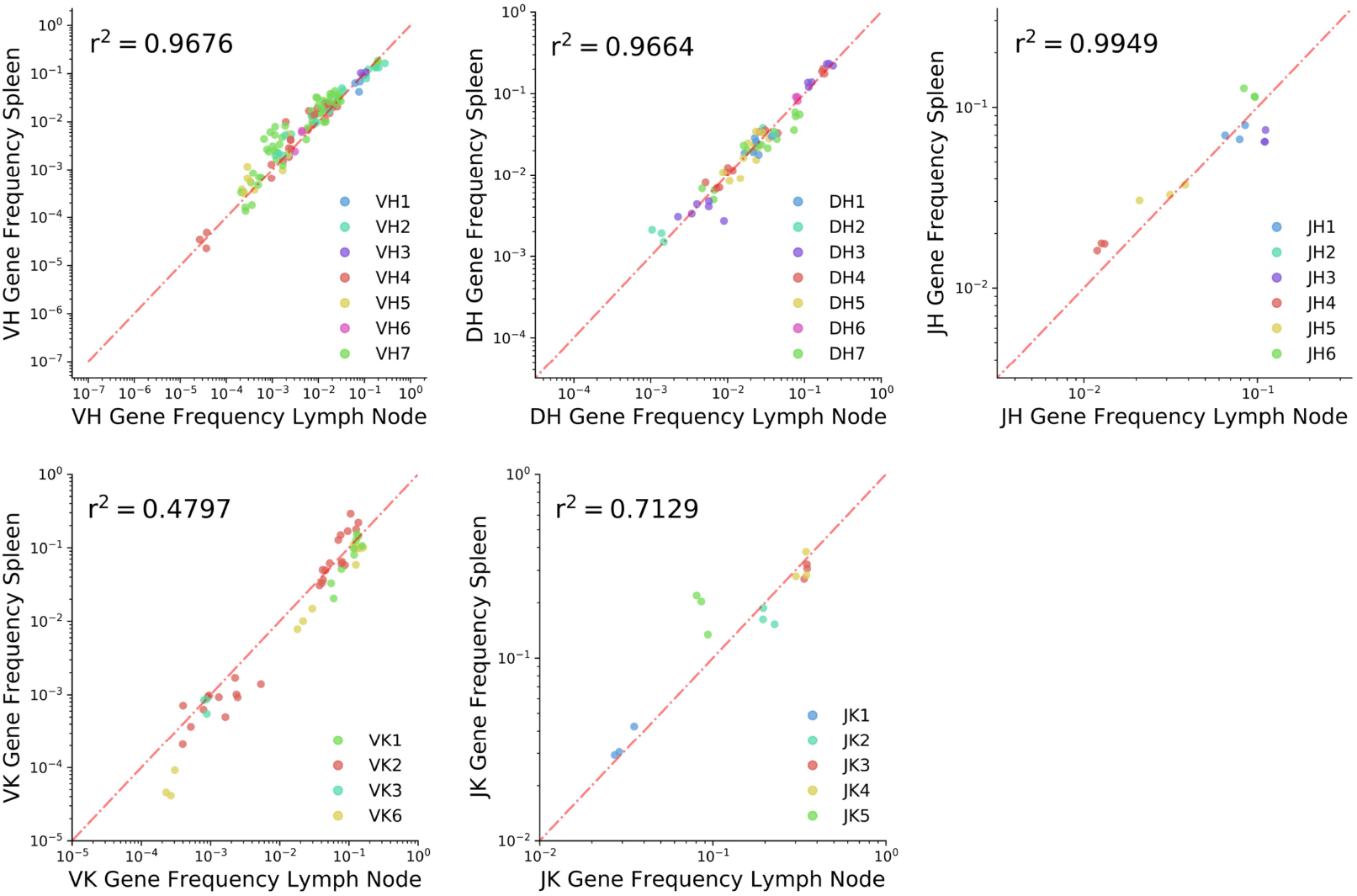
Intra-animal gene segment usage frequency scatter plots.

**Supplementary Figure 2.**
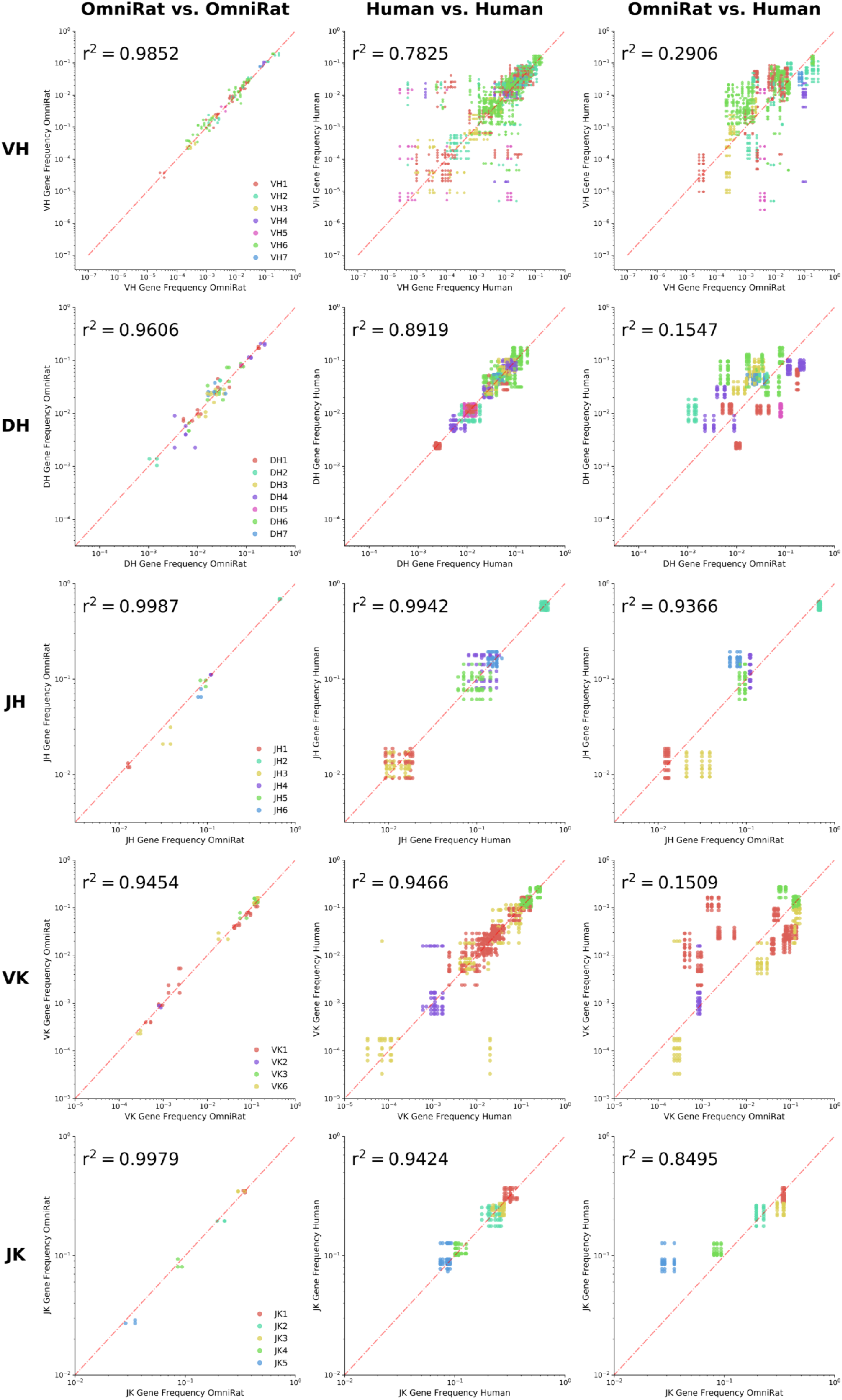
Intra-species and inter-species gene segment usage frequency scatter plots.

**Supplementary Figure 3.**
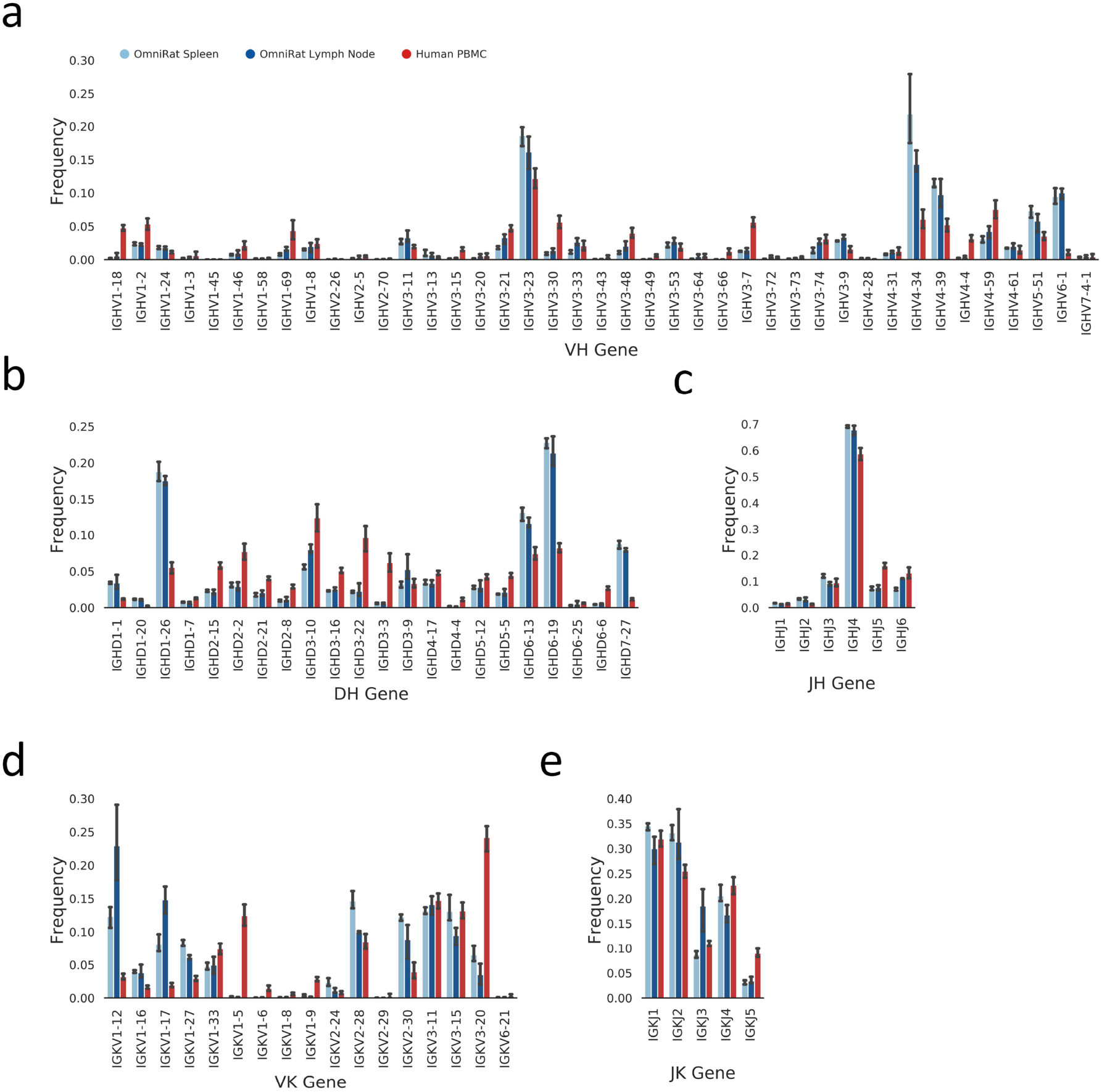
Gene segment usage segment frequencies. for a) VH, b) DH, c) JH, d) VK, e) JK. Species are colored for all figures as in (a)

**Supplementary Figure 4.**
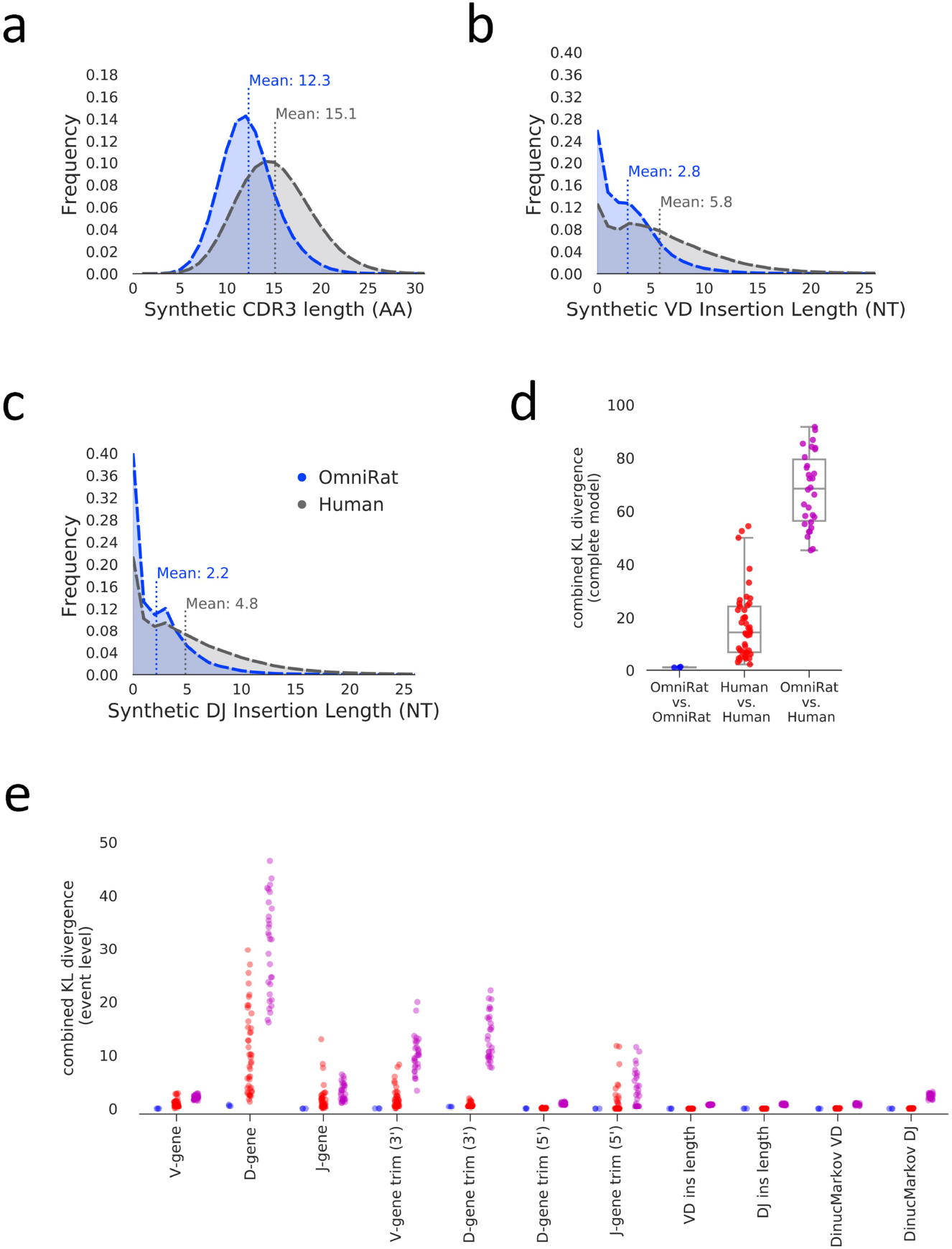
Naïve Repertoire modeling with IGoR. a) Synthetic CDRH3 length distribution for each species. Species are colored as in (c). CDRH3 lengths were determined using the ImMunoGeneTics (IMGT) numbering scheme. b) Synthetic VD insertion length distributions for each species. Species are colored as in (c). c) Synthetic DJ insertion length distributions for each species. Species are colored as in (c). d) Combined Kullback–Leibler divergence (KL divergence) between pairs of OmniRat models (blue), between pairs of human models (red), or between pairs of OmniRat and human models (purple).

**Supplementary Figure 5.**
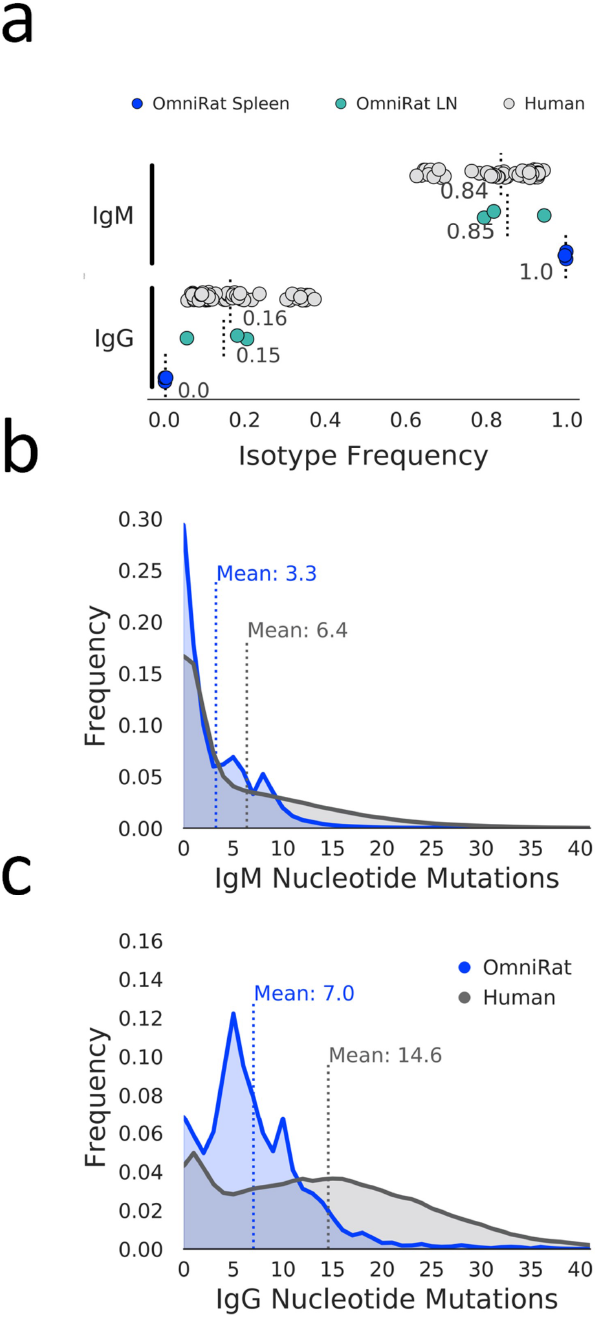
Class-switch recombination and somatic hypermutation. a) Sequence frequency by antibody isotype. b) Variable heavy gene nucleotide mutation number distributions for IgM sequences. Species colored as in (c). c) Variable heavy gene nucleotide mutation number distributions for IgG sequences.

**Supplementary Figure 6.**
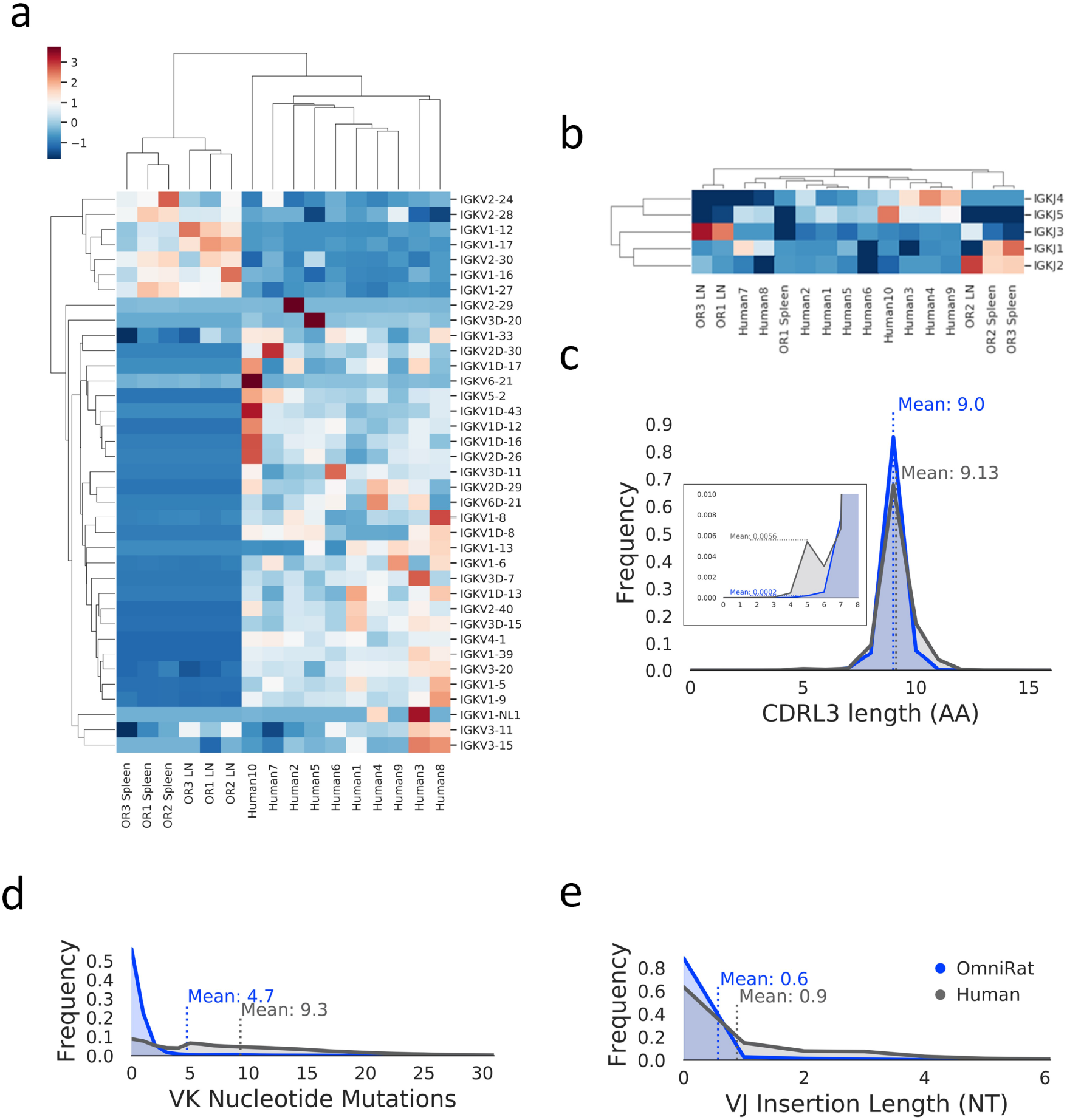
Kappa chain summary. Heatmaps of variable kappa (VK) gene usage in (a) and joining kappa (JK) gene usage in (b). Columns are antibody repertoires and rows are gene segments. Data was scaled by calculating theZ-score for each gene (row) and hierarchical clustering (Euclidean distance metric) was done. A dendrogram representation of clustering is shown. c) CDRL3 length distribution for each species. Species are colored as in (e). d) Variable kappa gene nucleotide mutation number distributions. e) VJ insertion length distributions for each species.

**Supplementary Table 1.**
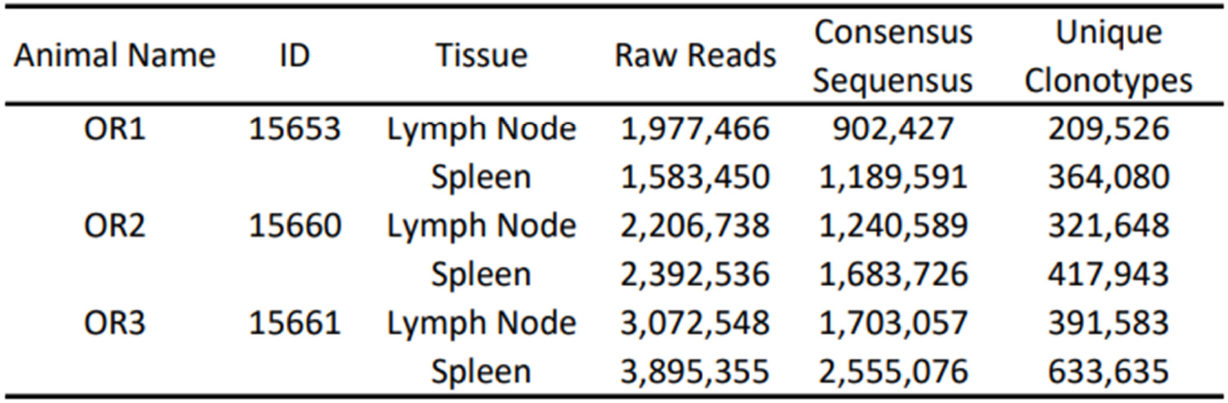
OmniRat heavy chain sequencing summary.

| Animal Name | ID | Tissue | Raw Reads | Consensus Sequences | Unique Clonotypes |
| --- | --- | --- | --- | --- | --- |
| OR1 | 15653 | Lymph Node | 1,977,466 | 902,427 | 209,526 |
|  |  | Spleen | 1,583,450 | 1,189,591 | 364,080 |
| OR2 | 15660 | Lymph Node | 2,206,738 | 1,240,589 | 321,648 |
|  |  | Spleen | 2,392,536 | 1,683,726 | 417,943 |
| OR3 | 15661 | Lymph Node | 3,072,548 | 1,703,057 | 391,583 |
|  |  | Spleen | 3,895,355 | 2,555,076 | 633,635 |

**Supplementary Table 2.**
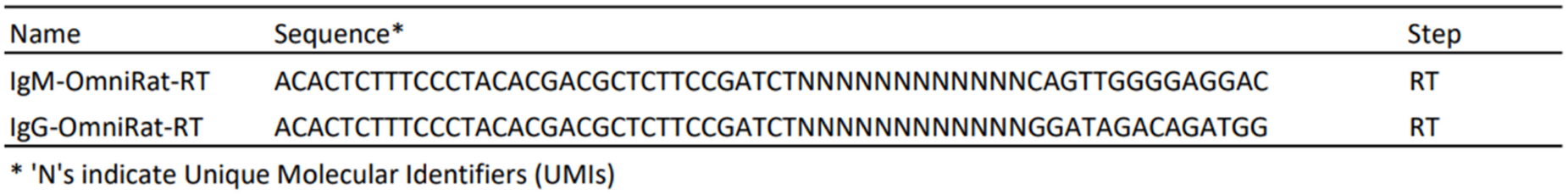
OmniRat heavy chain reverse transcription primers.

## References

1. Hu, J. K. et al. Murine antibody responses to cleaved soluble HIV-1 envelope trimers are highly restricted in specificity. J. Virol. 89, 10383–10398 (2015).

2. Jardine, J. et al. Rational HIV immunogen design to target specific germline B cell receptors. Science 340, 711–716 (2013).

3. Sok, D. et al. Priming HIV-1 broadly neutralizing antibody precursors in human ig loci transgenic mice. Science 353, 1557–1560 (2016).

4. Pantophlet, R. et al. Bacterially derived synthetic mimetics of mammalian oligomannose prime antibody responses that neutralize HIV infectivity. Nat. Commun. 8, 1601 (2017).

5. Brüggemann, M. et al. A repertoire of monoclonal antibodies with human heavy chains from transgenic mice. Proc. Natl. Acad. Sci. U. S. A. 86, 6709–6713 (1989).

6. Green, L. L. et al. Antigen-specific human monoclonal antibodies from mice engineered with human ig heavy and light chain YACs. Nat. Genet. 7, 13–21 (1994).

7. Lonberg, N. et al. Antigen-specific human antibodies from mice comprising four distinct genetic modifications. Nature 368, 856–859 (1994).

8. Brüggemann, M. et al. Human antibody production in transgenic animals. Arch. Immunol. Ther. Exp. 63, 101–108 (2015).

9. Green, L. L. Transgenic mouse strains as platforms for the successful discovery and development of human therapeutic monoclonal antibodies. Curr. Drug Discov. Technol. 11, 74–84 (2014).

10. Nemazee, D. Mechanisms of central tolerance for B cells. Nat. Rev. Immunol. 17, 281–294 (2017).

11. Osborn, M. J. et al. High-affinity IgG antibodies develop naturally in ig-knockout rats carrying germline human IgH/Ig*κ*/Ig*λ* loci bearing the rat CH region. J. Immunol. 190, 1481–1490 (2013).

12. Ma, B. et al. Human antibody expression in transgenic rats: comparison of chimeric IgH loci with human VH, D and JH but bearing different rat c-gene regions. J. Immunol. Methods 400-401, 78–86 (2013).

13. Briney, B., Inderbitzin, A., Joyce, C. & Burton, D. R. Commonality despite exceptional diversity in the baseline human antibody repertoire. Nature 566, 393–397 (2019).

14. Briney, B., Le, K., Zhu, J. & Burton, D. R. Clonify: unseeded antibody lineage assignment from next-generation sequencing data. Sci. Rep. 6, 23901 (2016).

15. Vollmers, C., Sit, R. V., Weinstein, J. A., Dekker, C. L. & Quake, S. R. Genetic measurement of memory b-cell recall using antibody repertoire sequencing. Proc. Natl. Acad. Sci. U. S. A. 110, 13463–13468 (2013).

16. He, L. et al. Toward a more accurate view of human b-cell repertoire by next-generation sequencing, unbiased repertoire capture and single-molecule barcoding. Sci. Rep. 4, 6778 (2014).

17. Briney, B. & Burton, D. R. Massively scalable genetic analysis of antibody repertoires (2018). Preprint available at https://www.biorxiv.org/content/early/2018/10/19/447813.

18. Umotoy, J. et al. Rapid and focused maturation of a VRC01-Class HIV broadly neutralizing antibody lineage involves both binding and accommodation of the N276-Glycan. Immunity 51, 141–154.e6 (2019).

19. Jardine, J. G. et al. HIV-1 VACCINES. priming a broadly neutralizing antibody response to HIV-1 using a germline-targeting immunogen. Science 349, 156–161 (2015).

20. Andrabi, R. et al. Identification of common features in prototype broadly neutralizing antibodies to HIV envelope V2 apex to facilitate vaccine design. Immunity 43, 959–973 (2015).

21. Marcou, Q., Mora, T. & Walczak, A. M. High-throughput immune repertoire analysis with IGoR. Nat. Commun. 9, 561 (2018).

22. Lee, E.-C. et al. Complete humanization of the mouse immunoglobulin loci enables efficient therapeutic antibody discovery. Nat. Biotechnol. 32, 356–363 (2014).

23. Longo, N. S., Rogosch, T., Zemlin, M., Zouali, M. & Lipsky, P. E. Mechanisms that shape human antibody repertoire development in mice transgenic for human ig H and L chain loci. J. Immunol. 198, 3963–3977 (2017).

24. Briney, B. S., Willis, J. R., Finn, J. A., McKinney, B. A. & Crowe, J. E., Jr. Tissue-specific expressed antibody variable gene repertoires. PLoS One 9, e100839 (2014).

25. Hoi, K. H. & Ippolito, G. C. Intrinsic bias and public rearrangements in the human immunoglobulin V*λ* light chain repertoire. Genes Immun. 14, 271–276 (2013).

26. Collins, A. M. & Watson, C. T. Immunoglobulin light chain gene rearrangements, receptor editing and the development of a Self-Tolerant antibody repertoire. Front. Immunol. 9, 2249 (2018).

27. Soto, C. et al. High frequency of shared clonotypes in human B cell receptor repertoires. Nature 566, 398–402 (2019).

28. Love, M. I., Huber, W. & Anders, S. Moderated estimation of fold change and dispersion for RNA-seq data with DESeq2. Genome Biol. 15, 550 (2014).

